# A generalized theoretical framework to investigate multicomponent actin dynamics

**DOI:** 10.1101/2024.12.10.627743

**Authors:** Mintu Nandi, Shashank Shekhar, Sandeep Choubey

## Abstract

The length of actin filaments is regulated by the combined action of hundreds of actin-binding proteins. While the roles of individual proteins are well understood, how they combine to regulate actin dynamics in vivo remains unclear. Recent advances in microscopy have enabled precise, high-throughput measurements of filament lengths over time. However, the absence of a unified theoretical framework has hindered a mechanistic understanding of the multicomponent regulation of actin dynamics. To address this, we propose a general kinetic model that captures the combined effects of multiple regulatory proteins on actin dynamics. We provide closed-form expressions for both time-dependent and steady-state moments of the filament length distribution. Our framework not only differentiates between various regulatory mechanisms but also serves as a powerful tool for interpreting current data and driving future experiments.

## I. INTRODUCTION

Actin filaments are key components of the cytoskeleton, driving processes such as cell motility, cytokinesis, endocytosis, and wound healing [1–3]. The regulation of filament length, essential for these processes, is controlled by a variety of actin-binding proteins (ABPs) that promote elongation, depolymerization, or capping [4–9]. While decades of biochemical research have helped elucidate the individual effects of many of these proteins, how their activities combine to drive complex actin dynamics in vivo remains poorly understood.

Recent advances in fluorescence microscopy have enabled the precise measurement of filament length changes over time for hundreds of filaments, producing rich datasets on the distribution of filament lengths as a function of time [10–13]. While numerous experimental studies have provided a wealth of data, the lack of a “theory of the experiment” [14] has hindered efforts to uncover the governing principles underlying multicomponent regulation of actin filaments.

To address this gap, we propose a general theoretical framework that captures the simultaneous effects of multiple ABPs on actin dynamics. We focus on two key aspects: (1) the time evolution of filament length distributions and (2) the steady-state distribution of filament lengths within a fixed time window. We derive exact closed-form expressions for the moments of filament length. To demonstrate the utility of this framework, we model the effects of two regulatory proteins with distinct mechanisms—an elongator (such as formin [15]) and a capper (such as capping protein [16]), both of which have been shown to simultaneously bind filament ends[17, 18]. We show that the mean and variance of filament length changes provide a powerful means to discriminate between these mechanisms. Our framework not only enables the analysis of existing experimental data but also provides a guide for designing new experiments.

## II. MODEL

We develop a generalized kinetic model in which individual actin filaments can stochastically transition between different states defined by the presence of specific ABPs on filaments. In each state, filaments can polymerize, or depolymerize, or remain capped. Our model assumes that the filament can exist in a total of *N* distinct states, with *N*_1_ states leading to polymerization and *N*_2_ states leading to depolymerization. We consider the capped states to be part of the first cohort without any loss of generality. The rate of polymerization or depolymerization in the *i*-th state is given by *r*_*i*_, and the transition rate from the *j*-th state to the *i*-th state is denoted by *k*_*ij*_. Our model involves two random variables: the state of the filament *i*, and the change in its length, Δ*L*_*t*_, over a given time interval (*t*). The change in length is defined as Δ*L*_*t*_ = *L*_*t*_ − *L*_0_, where *L*_*t*_ is the filament’s length at time *t* and *L*_0_ is its initial length at *t* = 0. Δ*L*_*t*_ can be positive (polymerization) or negative (depolymerization). We seek to compute the probability *P*_*i*_(Δ*L*_*t*_) that the filament’s length changes by Δ*L*_*t*_ over a time interval *t*, while in the *i*-th state. The master equation for the time evolution of *P*_*i*_(Δ*L*_*t*_)s in matrix form is given by

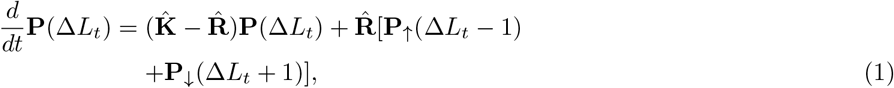

where the probability vector **P**(Δ*L*_*t*_) = (*P*_1_(Δ*L*_*t*_), …, *P*_*N*_ (Δ*L*_*t*_), *P*_*N* +1_(Δ*L*_*t*_), …, *P*_*N*_ (Δ*L*_*t*_))^*T*^.

To obtain analytical solutions of Eq. (1), we introduce separate probability vectors for polymerization and depolymerization (Fig. S1). Specifically, **P**_↑_(Δ*L*_*t*_ − 1) represents the probabilities of polymerizing states, and **P**_↓_(Δ*L*_*t*_ + 1) represents the probabilities of depolymerizing states. The elements in **P**_↑_ are zero for depolymerizing states, and vice versa for **P**_↓_ (Fig. S1). The matrix 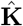 in the master equation (1) describes state transitions, with off-diagonal elements *k*_*ij*_ representing the rate of transition from state *j* to *i*, and diagonal elements *k*_*ii*_ representing the outflow rate from state *i* (Fig. S1). The matrix 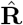 is diagonal, representing the rates of polymerization and depolymerization (Fig. S1). From Eq. (1), we derive exact expressions for the moments of 1) the distribution of Δ*L*_*t*_ over time, and 2) the steady-state distribution of Δ*L*_*τ*_ within a fixed time window *τ*, where steady-state refers to the long-time limit where the probability distribution of Δ*L*_*τ*_ ceases to change.

We calculate the *n*th moment of the distribution of Δ*L*_*t*_ by multiplying both sides of Eq. (1) by 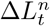. By summing over all values of Δ*L*_*t*_ and finally multiplying both sides by 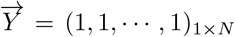, as described in the *SI text*, we obtain the following equation for the *n*-th moment,

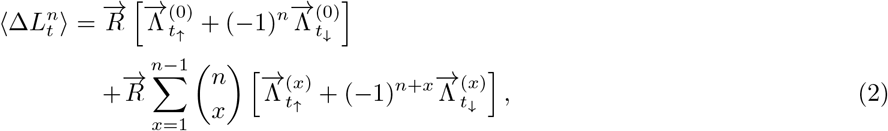

where, 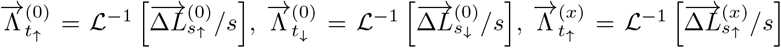, and 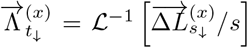 (see *SI text* for details). Here, ℒ^−1^ denotes the inverse Laplace transformation of the partial moment vectors 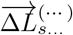 defined in Laplace space *s*. We note here that 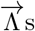 are obtained as functions of the matrices 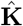, and 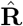 (see *SI text*).

Next, we calculate the *n*-th moment of the steady-state distribution of Δ*L*_*τ*_, given by

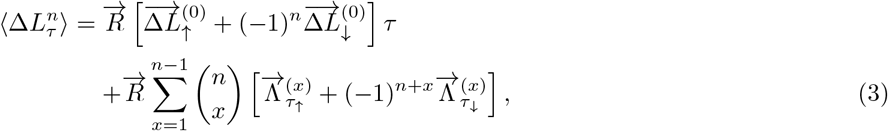

where, 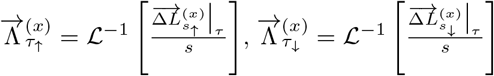 (see *SI text* for detailed analytical steps). Here, 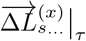 denotes the steady-state partial moment vectors. In the above equation, 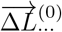 stands for the zeroth order partial moment vectors, which characterize the occupancy of various states. The 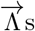 are obtained as functions of the the matrices 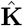, and 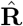.

Below, we utilize our analytical results to dissect specific regulatory mechanisms of actin dynamics.

## III. REGULATION OF ACTIN FILAMENT LENGTH BY A SINGLE ABP

To explore how filament growth distributions reveal mechanistic insights into multicomponent regulation of actin dynamics, we examine the combined effects of an elongator and a capper protein on filament length. Elongators promote growth [4, 5], while cappers inhibit polymerization[6]. We consider three scenarios: 1) actin filaments with only an elongator, 2) actin filaments with only a capper, and 3) actin filaments with both proteins.

### A. Effect of an elongator on actin filament length

We first examined how an elongator affects actin filament length. In the presence of an elongator, the filament can exist in two states: a bare state (B) and an elongator-bound state (BF), with corresponding polymerization rates *r*_1_ and *r*_2_. The binding and unbinding rates of the elongator are 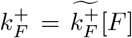 and 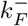, where [*F*] is the elongator concentration. The mean and variance of Δ*L*_*t*_, as a function of time and biochemical rates, are given by

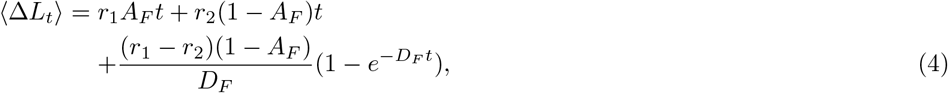

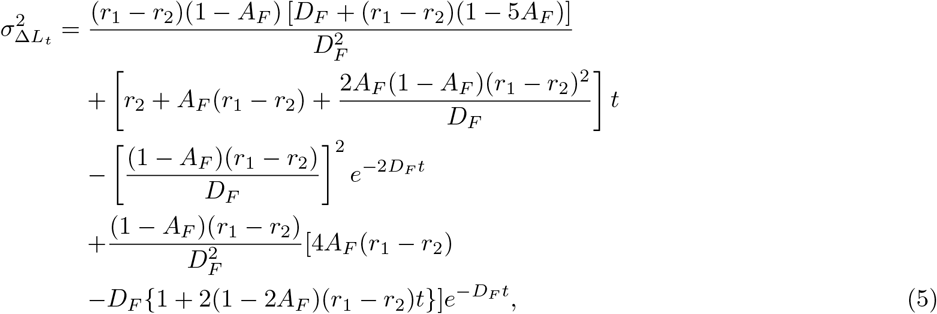

where 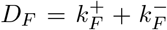 and 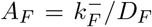. When [*F*] = 0, actin filament length increases linearly over time, driven solely by the polymerization rate of the bare (B) state (Fig. 2B). With elongator present, growth remains linear initially but becomes nonlinear at intermediate times due to the transition to the BF state. In the long term, growth is governed by the BF state polymerization rate. The Fano factor (variance over mean), which quantifies growth variability, remains at one when [*F*] = 0. In the presence of elongator, it rises above one, peaks, and then decreases, approaching one (Fig. 2B). The intermediate increase in the Fano factor reflects slow switching between the B and BF states, driven by elongator concentration.

**Figure 1.**
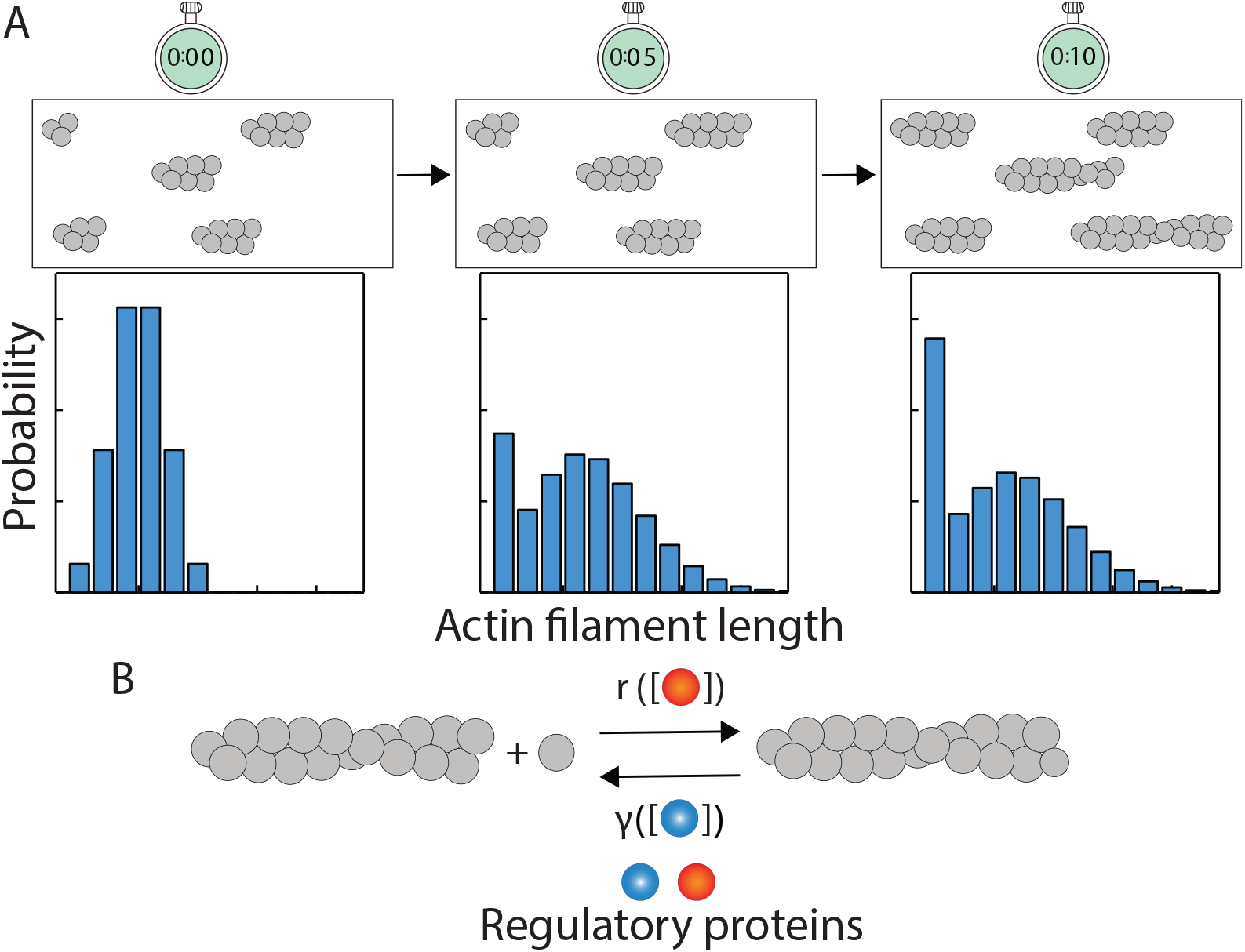
Time evolution of actin filament length. (A) Experimental time traces of actin filament length offer snapshots of the length distribution at successive time points [19–21]. (B) Effect of different regulatory proteins on actin dynamics.

**Figure 2.**
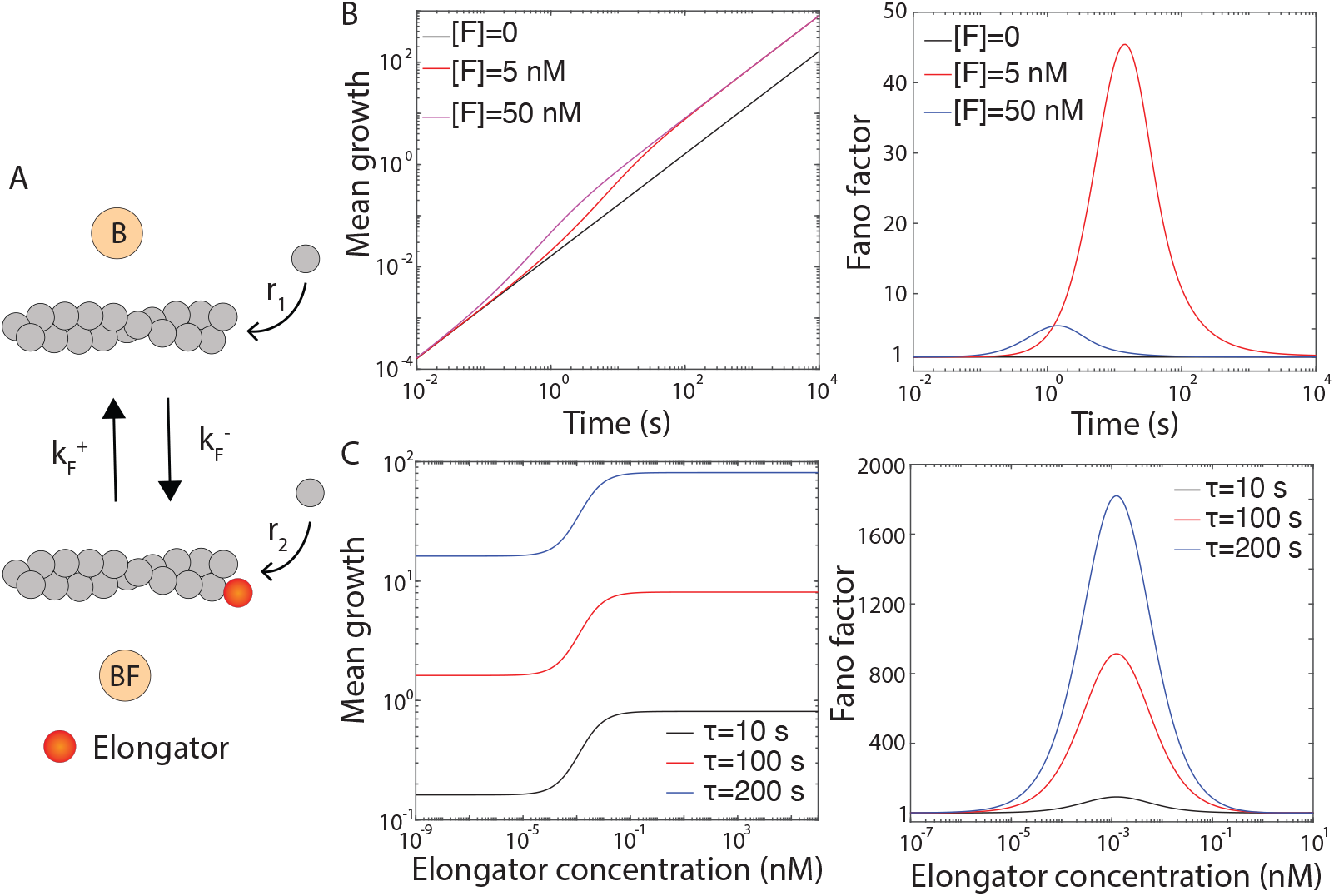
Effect of an elongator on actin filament length. (A) Two-state model of actin polymerization in the presence of elongator is illustrated. (B) Mean filament growth (*µm*) and Fano factor are plotted as a function of time for varying elongator concentrations. (C) Steady-state mean growth (*µm*) and Fano factor are shown as functions of elongator concentration for different *τ* values. Following parameters were used for elongator 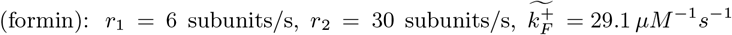, and 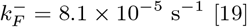.

The closed-form expressions for the steady-state mean and variance of Δ*L*_*τ*_ within a fixed time window *τ*, are given by

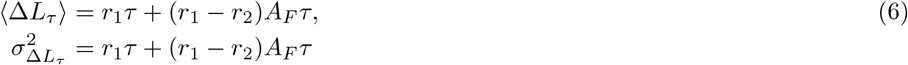

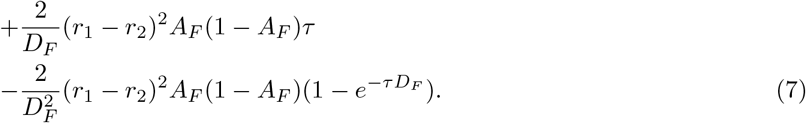

At steady-state, mean filament growth increases with elongator concentration, plateauing at high concentrations (Fig. 2C). The Fano factor shows nonmonotonic behavior: it remains at one at low concentrations, rises above one, peaks, and then decreases, approaching one at high concentrations (Fig. 2C). The increase at intermediate concentrations is due to slow switching between the B and BF states.

### B. Effect of a capper on actin filament length

Next, we examine how a capper protein affects actin filament length. The filament can exist in two states: a bare state (B) and a capper-bound state (BC), with polymerization rates *r*_1_ and *r*_2_ = 0, respectively. The capper binding and unbinding rates are 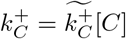 and 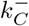. The mean and variance of the distribution of Δ*L*_*t*_ are given by

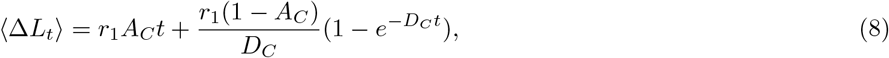

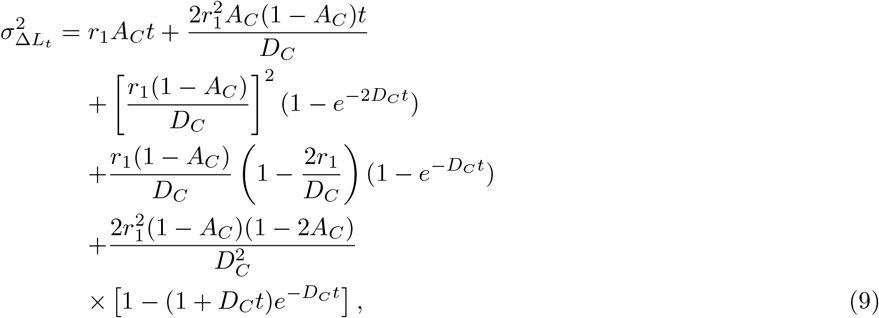

where 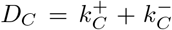 and 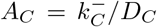. At [*C*] = 0, the mean growth is higher than when [*C*] *>* 0 (Fig. 3B). As capper concentration increases, the growth rate decreases, demonstrating an inverse relationship between [*C*] and filament growth (Fig. 3B).

**Figure 3.**
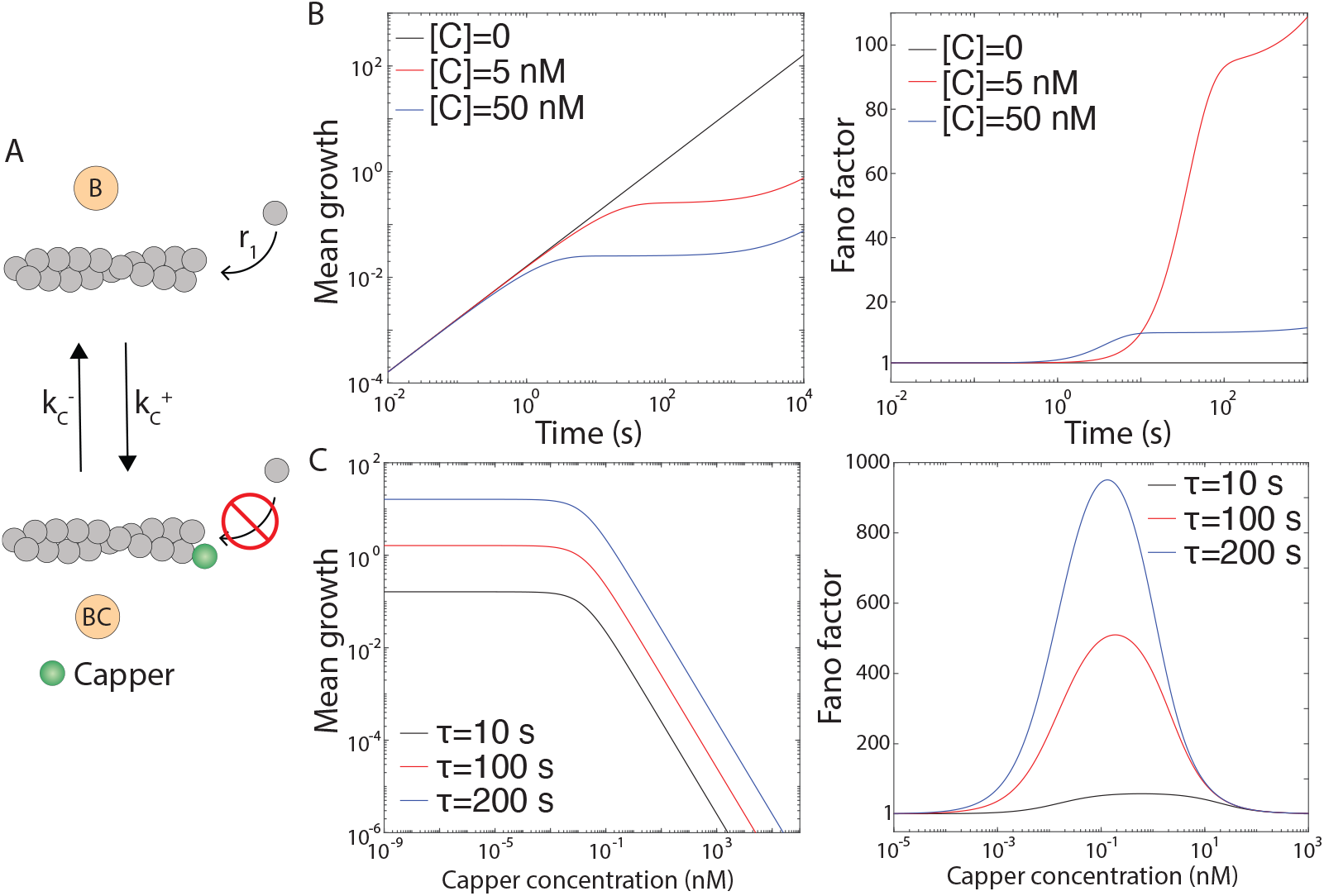
Effect of a capper on actin filament length. (A) Two-state model of actin dynamics in the presence of a capper is shown. (B) Mean filament growth (*µm*) and Fano factor are plotted as functions of time for different capper concentrations. (C) Steady-state mean growth (*µm*) and Fano factor are shown as functions of capper concentration for different values of *τ*. Following parameters were used for the capper(Capping protein):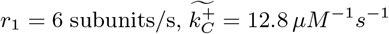, and 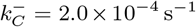 [19].

When [*C*] = 0, the Fano factor is one, and filament growth is Poissonian (Fig. 3B). Adding capper initially has no effect on growth variability, as it does not bind to the barbed end. Over time, capper binds, capping the filament and increasing the Fano factor, with a more pronounced increase at lower concentrations. The transient phase exhibits dynamic behavior due to frequent transitions between the bare (B) and capped (BC) states. In the long term, the Fano factor plateaus (Fig. 3B).

The mean and the variance of the steady-state distribution of Δ*L*_*τ*_ are given by

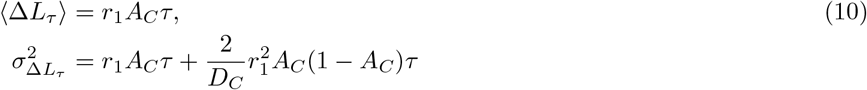

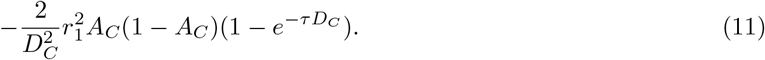

Mean steady-state growth is highest without capper and decreases with increasing capper concentration (Fig. 3C), as capper reduces the time filaments spend in the bare (B) state, where polymerization occurs. The Fano factor remains at one at low capper concentrations but shows non-monotonic behavior: it rises, peaks when B and BC states are equally occupied, and then returns to one (Fig. 3C). Over longer time windows, variability increases due to higher overall growth.

## IV. COMBINED EFFECTS OF AN ELONGATOR AND A CAPPER ON ACTIN FILAMENT LENGTH

So far we have quantified the individual effects of an elongator or a capper. However, in cells, these factors act simultaneously, often targeting the same site on a filament[17, 18]. We consider two models to explore the combined effect of an elongator and a capper, see Fig. 4A. First, Competitive Binding Model: The elongator and capper bind the same filament end in a mutually exclusive manner. In this model, the filament can exist in three states—free (B), capper-bound (BC), or elongator-bound (BF)—with polymerization rates *r*_1_, *r*_2_, and *r*_3_ = 0, respectively. Second, Simultaneous Binding Model: Both proteins can simultaneously bind the same filament end. Thus, the filament can now occupy four states—free (B, *r*_1_), elongator-bound (BF, *r*_2_), capper-bound (BC, *r*_4_ = 0), or dual-bound (BFC, *r*_3_ = 0).

**Figure 4.**
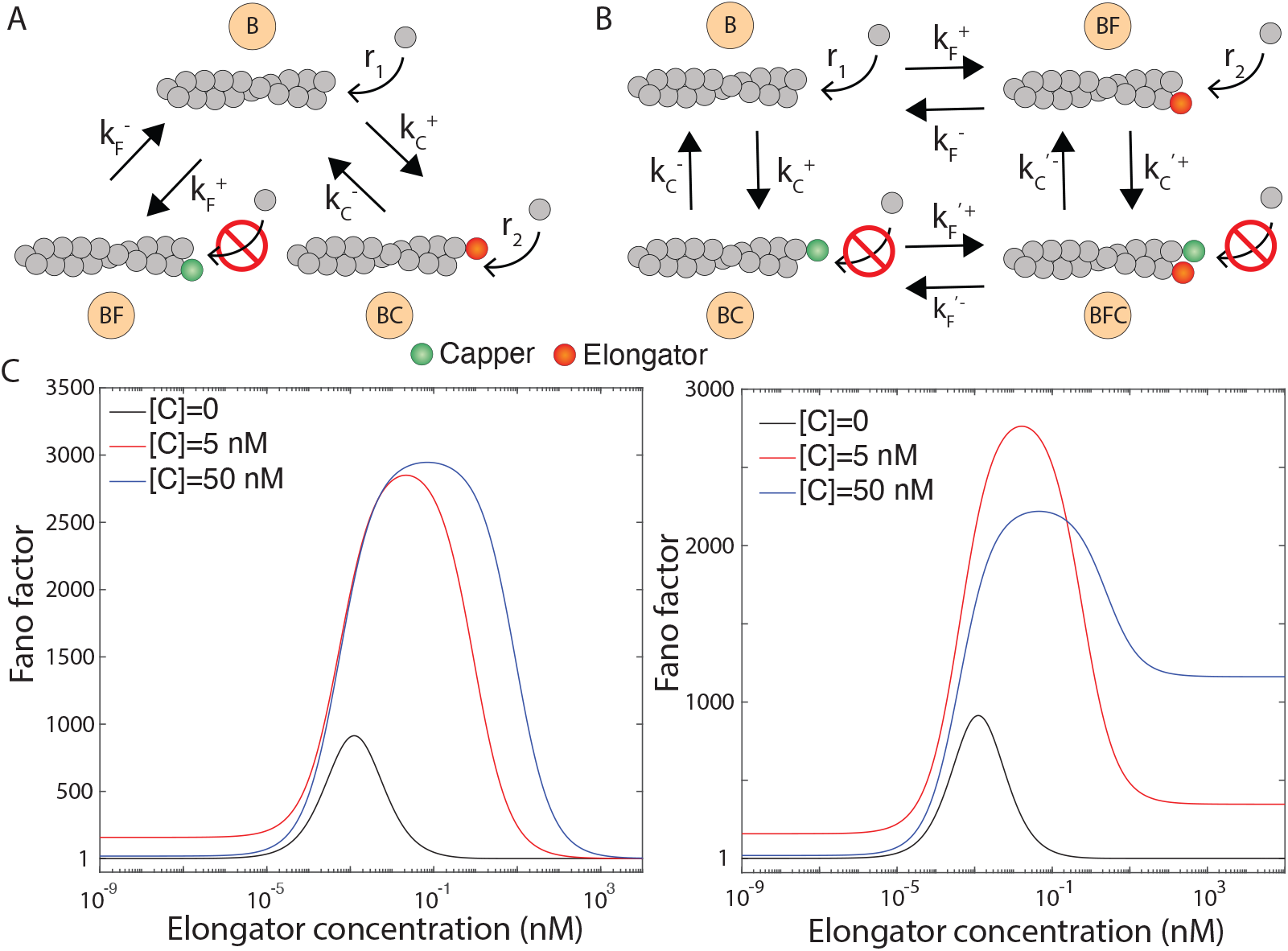
Combined effect of a capper and an elongator on actin filament length. (A) Competitive and (B) simultaneous binding models are shown. (C) Fano factor in steady-state is shown as a function of elongator concentration for various capper concentrations. Steady-state plots are generated for a fixed time window, *τ* = 100*s*. Following parameters for elongator (formin) and capper (capping protein) are used: 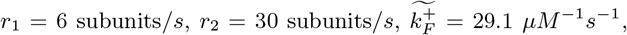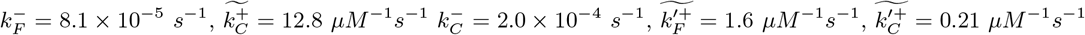, and 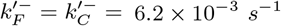 [19].

To differentiate between the two models, we analyze steady-state variability in filament growth by varying elongator concentration at different capper concentrations (Fig. 4B). In both models, the Fano factor shows non-monotonic behavior: it rises from low values at low elongator concentrations, peaks, and then decreases at higher concentrations. In the competitive binding model, the Fano factor approaches one at high elongator concentrations, indicating Poissonian growth. In contrast, the simultaneous binding model yields a Fano factor greater than one, which increases with capper concentration. These differences reflect how state occupancy changes with elongator concentration. In the competitive model, high elongator concentrations increase occupancy of the elongator-bound state, reducing variability. In the simultaneous model, the filament alternates between two states, leading to higher variability even at high elongator concentrations.

Different mechanisms of multicomponent regulation of actin dynamics can be discriminated by the distributions of filament lengths they produce.

## V. DISCUSSION

The study of actin filament dynamics has a rich experimental history, with theoretical models playing a key role in interpreting data. Two main classes of modeling efforts have emerged in this field. The first focuses on how actin filament length is regulated in steady-state conditions [22–25]. The second class of models examines the temporal aspects of filament growth [26, 27]. Notably, most of these models are computational and focus on specific mechanisms with a limited set of actin-binding proteins. In contrast, we propose a generalized analytical framework that captures the combined effects of an arbitrary number of regulatory proteins on actin filament dynamics.

So far, we have focused primarily on a single elongator and capper. However, cells contain a diverse range of regulatory factors, including depolymerases like twinfilin and cofilin [28], as well as numerous elongators and cappers such as formins [15] (with 15 mammalian isoforms) and Ena/VASP proteins [29]. Further complexity arises from the two distinct filament ends and the age of the filaments, which influence protein binding and activity. Due to its general framework, our model can accommodate these complexities, offering a more integrated understanding of actin dynamics in physiological contexts.

## ACKNOWLEDGMENTS

MN acknowledges SERB, India, for the National Post-Doctoral Fellowship [PDF/2022/001807]. SS was funded by NIH NIGMS Grant No. R35GM143050. SC acknowledges the support provided by the DBT Ramalingaswami Fellowship.

## Supplymentary material

### 1 Moments of actin filament length distribution in the presence of an arbitrary number of actin binding proteins

To compute all the moments of 1) filament length distributions as a function of time and 2) the steady-state distribution of filament lengths within a fixed time window, we use the master equation (1), as described in the main text. This equation can be exactly solved to obtain the *n*-th moment of the distribution of Δ*L*_*t*_ as a function of time and Δ*L*_*τ*_. To compute the transient moments, we define the following vectors of partial moments,

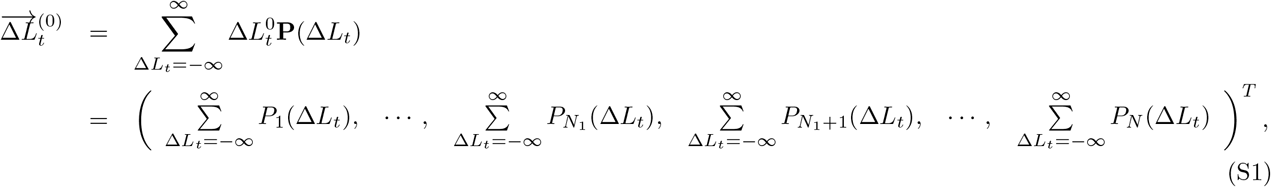

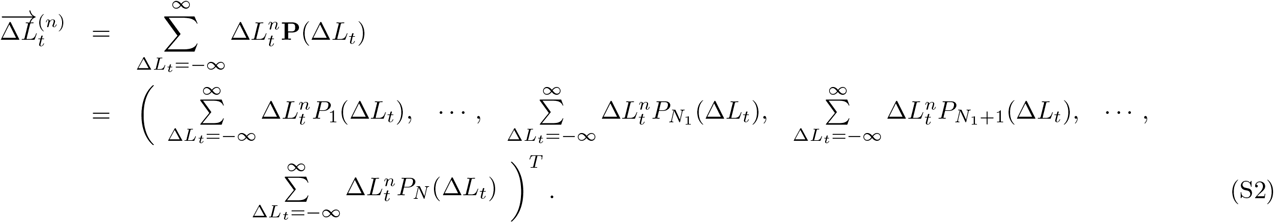

In Eq. (S1), 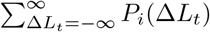 represents the probability of occurrence of filament state *i* (i.e., state visit probability). In Eq. (S2) 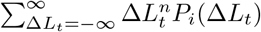 stands for the *n*^*th*^ partial moment due to the state *i*. These vectors are instrumental in calculating the moments of the probability distribution of the change in length of the filament. For instance, the *n*^*th*^ central moment can be expressed as the sum of all elements of the vector 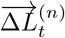, i.e.,

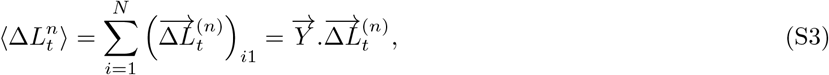

where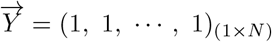. We note that 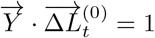, as the total state visit probability must always equal unity. To calculate 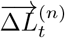, both sides of master equation (1) are multiplied by 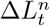, followed by summing over all values of Δ*L*_*t*_, which results in

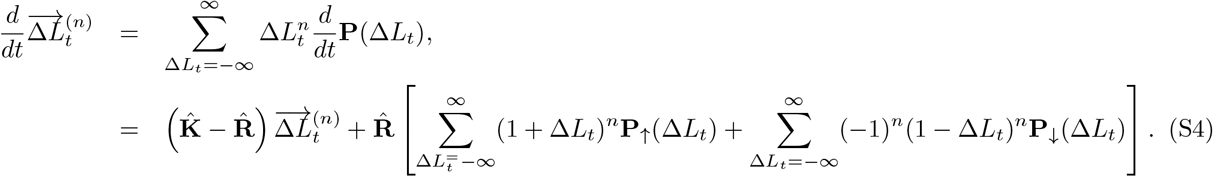

**Figure S1:**
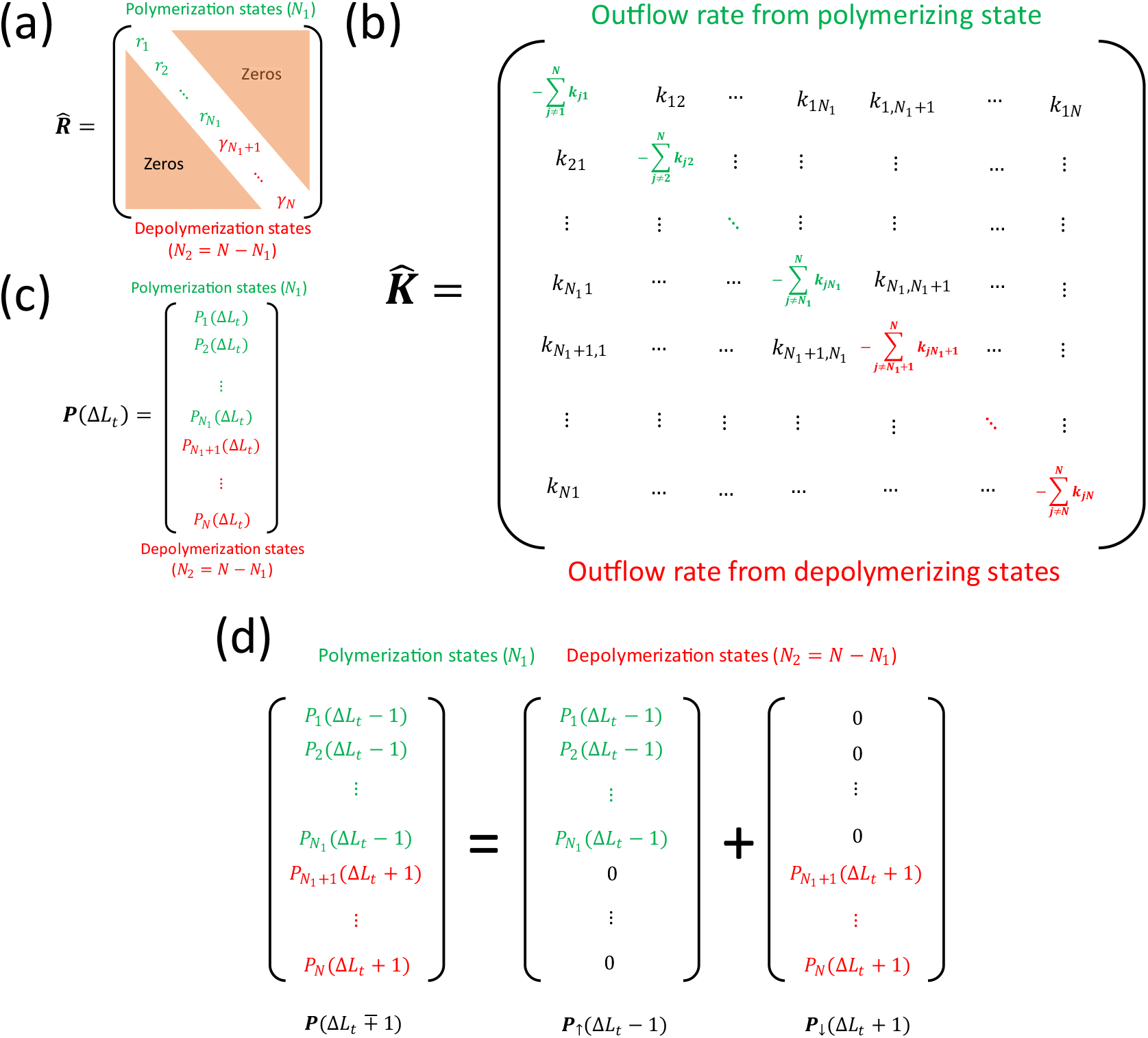
The schematics of structures of the matrices (a) 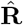, (b) 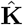, and (c) **P**(Δ*L*_*t*_). (d) The decomposition of transition probability vector.

In deriving Eq. (S4), we have employed the change of variable 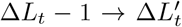 and 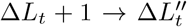 followed by revert back to Δ*L*_*t*_ for notational simplicity. We define the binomial expansions 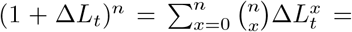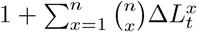 and 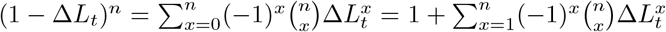, where 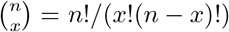 is called binomial coefficient. Employing the binomial expansion on Eq. (S4) yields,

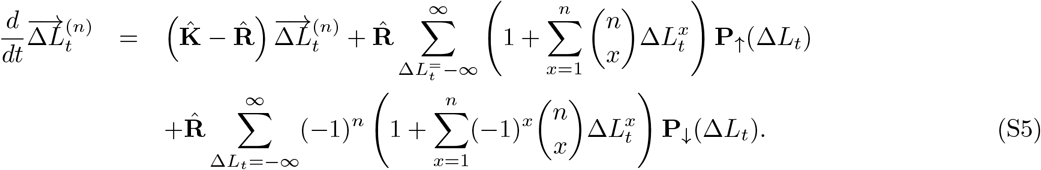

We now define 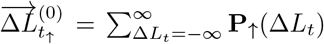 and 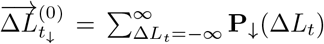 as the state visit probabilities from polymerizing and depolymerizing states, respectively. This gives 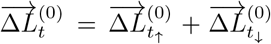 Again, the *n*^*th*^ partial moment vector follow the similar relation, i.e., 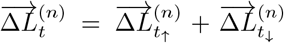 where partial moment vector due to polymerizing and depolymerizing states are defined as, 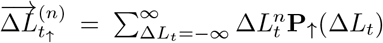 and 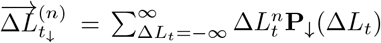, respectively. Using these definitions, we rearrange Eq. (S5) as,

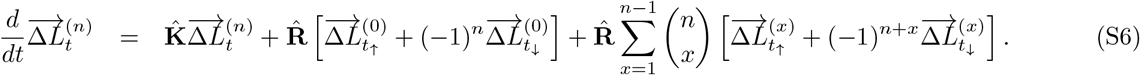

We multiply both sides of Eq. (S6) by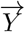 and use Eq. (S3), which leads to

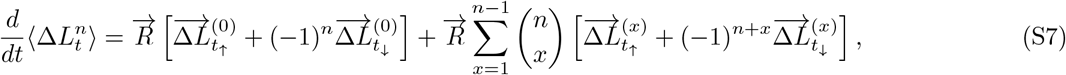

where, following the definition of matrices 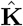 and 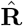, we arrive at 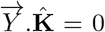 and 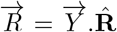 where 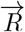 represents a row vector containing the diagonal elements of 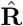. Eq. (S7) forms the *n*^*th*^ moment equation. Solving Eq. (S7) yields the expression of *n*^*th*^ central moment for Δ*L*_*t*_, provided we have the explicit expressions of 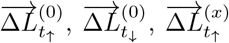, and 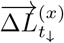. To solve this moment equation, we utilize the Laplace transformation with the initial condition that 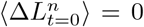, reflecting the fact that filament elongation or shortening has not started yet at initial times. This approach leads to the following expression

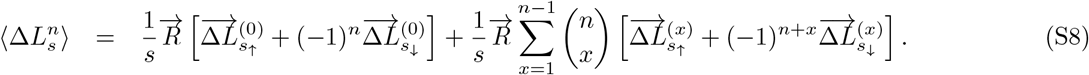

To find 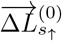 and 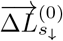 associated with Eq.(S8), we begin by computing the equation for 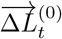 by summing master equation (1) over all values of Δ*L*_*t*_. This results in

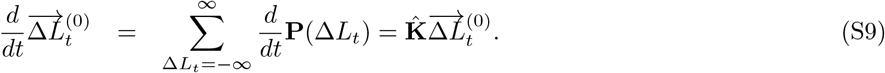

While deriving Eq. (S9), we use the change of variable 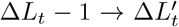 and 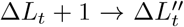. Solving Eq. (S9) in Laplace domain with the initial condition that at the initial time, the filament exists in state 1 (free state) which yields 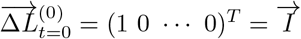. This operation results in,

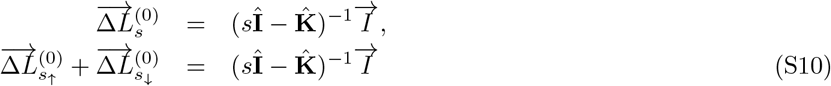

where 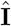 stands for the identity matrix. Now, it is important to decompose the vector obtained from the matrix operation of the right-hand side of Eq. (S10) and must correspond to the decomposition of **P**(Δ*L*_*t*_ ∓ 1) (see Fig. S1d). The decomposition of 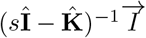 results in two vectors: one has non-zero values of the elements for polymerizing states and zero values for depolymerizing states, which provides the expression of 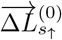. On the contrary, the second vector contains zero values of the elements for polymerizing states and non-zero values for depolymerizing states, resulting in 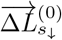. As the general matrix forms of 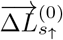 and 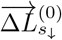 are difficult to show, the computation of these two vectors has been performed using suitable software.

To compute 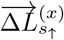 and 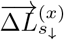, we derive the equation for 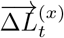 by rewriting Eq. (S6) in terms of *x*^*th*^ partial moment equation as,

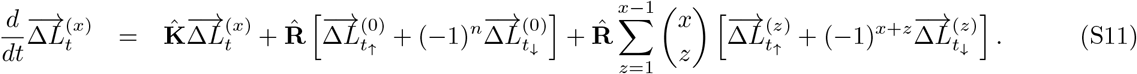

Upon Laplace transformation with the initial condition that at *t* = 0 the partial moments are zero, i.e., 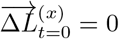, Eq. (S11) results in,

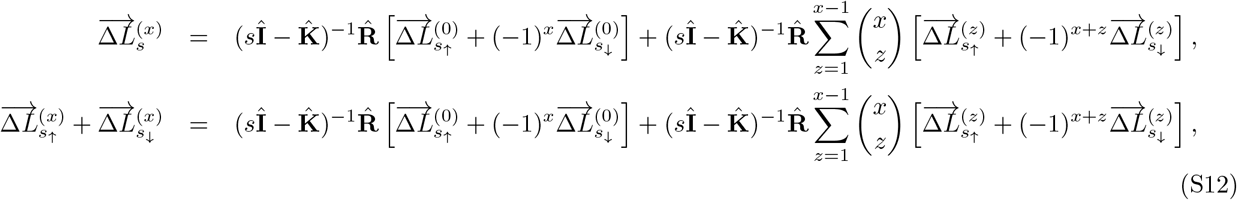

We know that 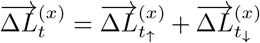 are also true in Laplace domain. Using this equality and following the previously stated decomposition of 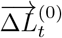, we compute the right-hand side of Eq. (S12), which yields a vector of dimension *N* ×1. We then decompose this vector into polymerizing and depolymerizing components: in the polymerizing vector, the first *N*_1_ elements are non-zero and the remaining *N*_2_ elements are zero; conversely, in the depolymerizing vector, the first *N*_1_ elements are set to zero and the next *N*_2_ elements are non-zero. This method of decomposition yields 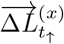 and 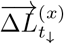.

Eq. (S8) together with Eqs. (S10) and (S12) compute the expression of *n*^*th*^ central moment in the Laplace domain. We write the expression of time-dependent *n*^*th*^ central moment of the change in length of the filament by performing inverse Laplace transformation on Eq. (S8), which gives

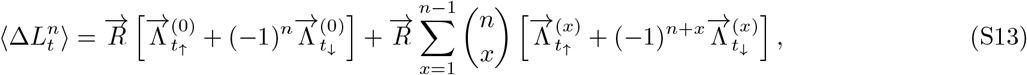

where, 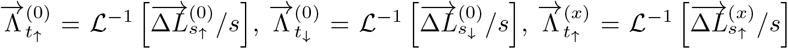, and 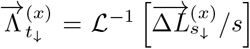.Here,the symbol ℒ^−1^ is used to refer the inverse Laplace transformation.

To compute the steady-state moments of actin length distribution, we begin by setting the left-hand side of Eq. (S9) to zero as the probabilities of visiting different states become independent of time and we denote it as 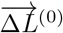. Additionally, the sum of state visit probabilities will be unity at steady-state. Mathematically, we express these two conditions as,

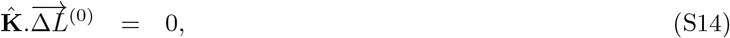

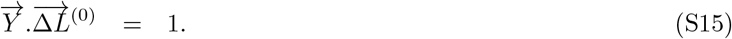

On solving these two coupled equations, the state visit probability vector 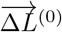 is computed. We decompose 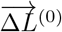 into two partial vectors: one is the polymerizing vector, 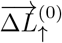, where the first *N*_1_ elements are non-zero and the remaining *N*_2_ elements are zero, and second the depolymerizing vector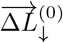, where the first *N*_1_ elements are zero and the next *N*_2_ elements are non-zero. We have used this method of decomposition in the case of deriving temporal moments.

We now replace the time variable *t* with *τ* in Eq. (S7) and perform Laplace transformation on both sides of the equation with the initial condition that 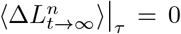. This initial condition is chosen to track the change in length of the filament that occurs within the window from *t* to *t* + *τ*, excluding any change in length that occurred up to time *t*. These operations lead to

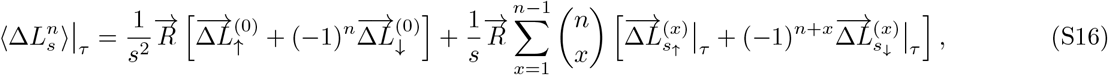

where, the terms 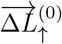 and 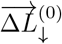 are known from Eqs. (S14-S15). The unknown quantities 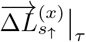 and 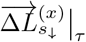 are computed by evaluating Eq. (S11) at steady-state and we then perform Laplace transformation with the initial condition that 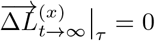. This results in,

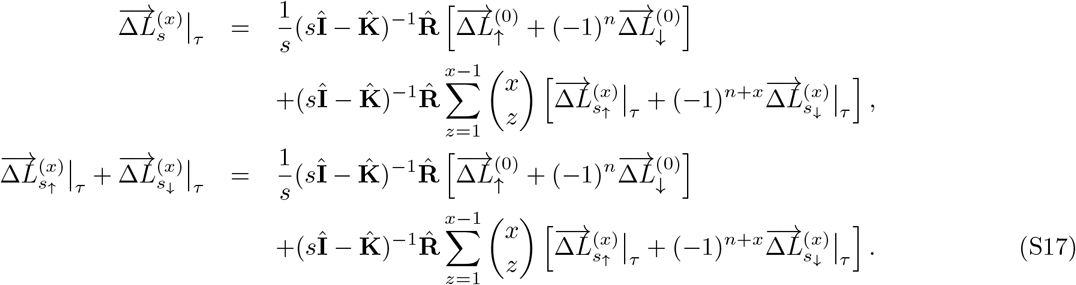

The decomposition of partial moment vector also applies to steady-state scenario. Using this decomposition, we have 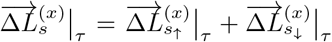. We now decompose the resultant vector obtained from the right-hand side of Eq. (S17), yielding two partial vectors: one corresponds to 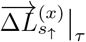 where the first *N*_*1*_ elements are non-zero but the next *N*_*2*_ elements are zero, and the second 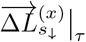, where the first *N*_*1*_ elements are zero but the next *N*_*2*_ elements are non-zero. Performing inverse Laplace transform of Eq. (S16) with the help of Eq. (S17), we have the *n*^*th*^ central moment as a function of time window *τ*,

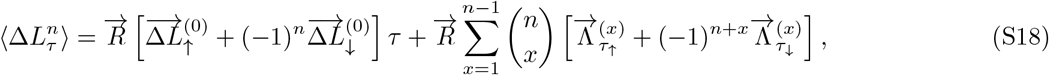

where, 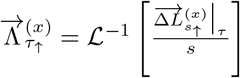 and 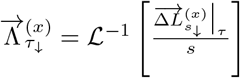.

Note that such master equation-based models have been employed in other fields of single-molecule biology to study their governing principles [1, 2, 3].

### 2 Moments of actin length distribution regulated by an elongator

In the presence of an elongator, the governing master equations are,

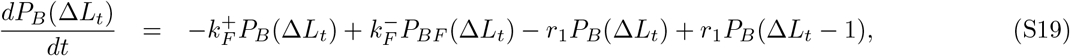

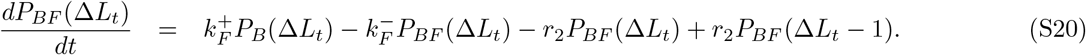

Expressing the above two equations in matrix form according to equation (1), we write the associated matrices as,

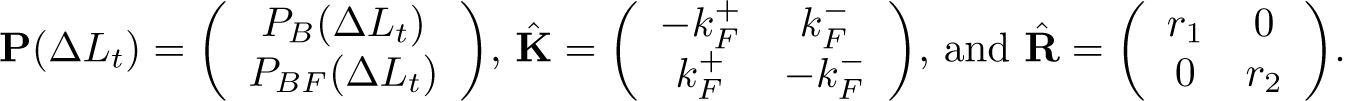

Using Eqs. (S13), we derive the corresponding moments (here we show up to the second moment) to obtain 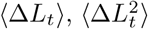. Using the first and second moments, we express the variance of 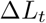 as 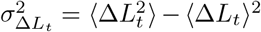. The closed-form analytical expressions for mean change in length ⟨Δ*L*_*t*_⟩, variance 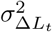 are given by

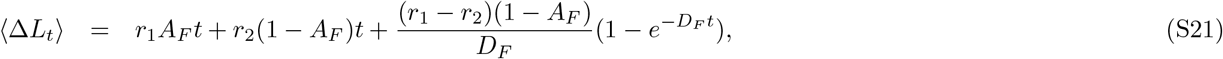

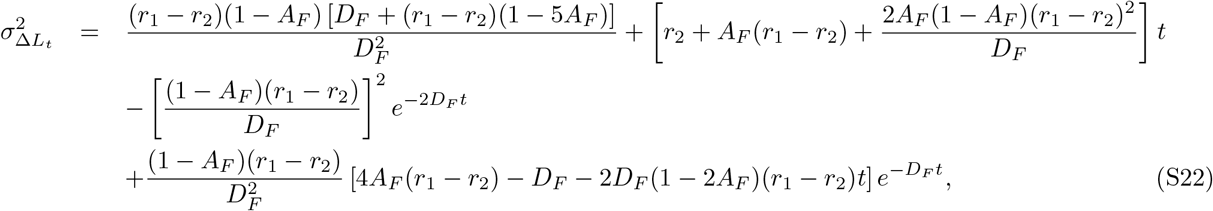

where 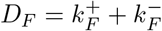 and 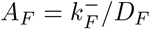.

Next, we compute the mean, variance at steady-state using Eq. (S18) and obtain 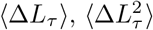. Using first and second central moments, we define the variance as 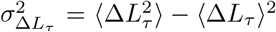. The closed-form expressions of mean and variance are written as

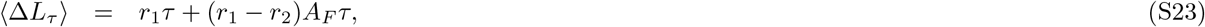

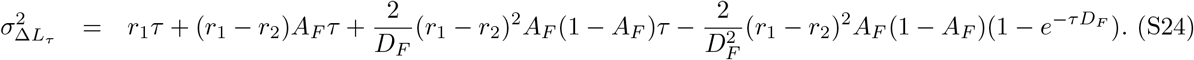

### 3 Moments of actin length distribution regulated by a capper

The governing master equations for filament length in the presence of a capper is given by

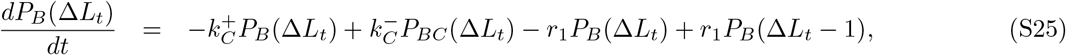

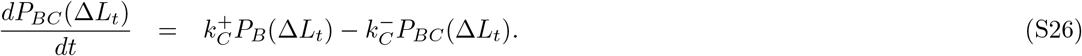

Expressing the above two equations in matrix form according to Eq. (**??**), the associated matrices are,

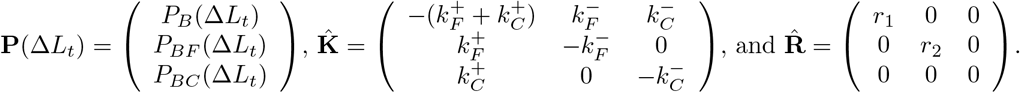

Utilizing the general solution provided in Eqs. (S13), we derive the corresponding moments (up to the second moment) to obtain ⟨Δ*L*_*t*_⟩, and 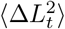. Using the first and second moments, we express the variance of Δ*L*_*t*_ as 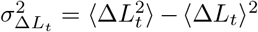 The closed-form analytical expressions for mean change in length ⟨Δ*L*_*t*_⟩, variance 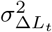 are given by,

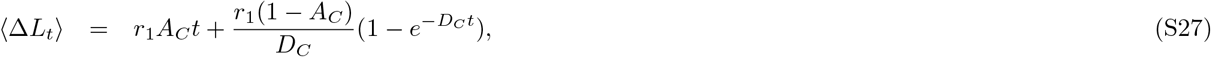

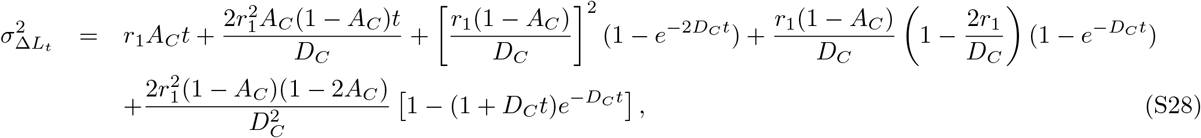

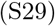

where 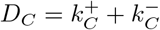 and 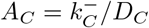.

Next, we compute the mean, variance, and third central moment at steady-state using Eq. (S18) and obtain ⟨Δ*L*_*τ*_ ⟩, and 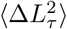. Using first and second central moments, we define the variance as 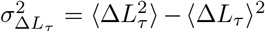. The closed-form expressions of mean and variance are written as,

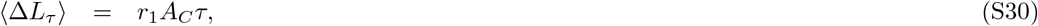

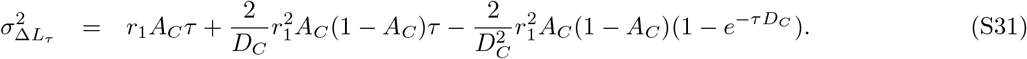

### 4 Moments of actin length distribution for competitive binding model and simultaneous binding models

#### 4.1 Competitive binding model

In this model, the filament undergoes elongation from states B and BF with rates *r*_1_ and *r*_2_, respectively. To write the master equation according to Eq. (1) described in the main text, we define the following matrices,

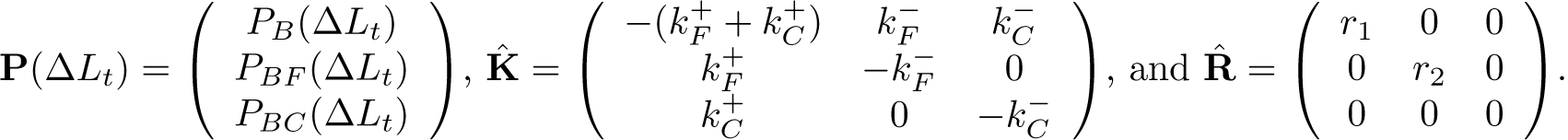

Using Eq. (S13), we derive the corresponding moments ⟨Δ*L*_*t*_⟩, and 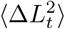. Using the first and second moments, the variance can be written as 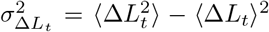. Due to the complexity of the analytical expressions, we don’t report them here.

Next, we calculate the mean, variance, and third central moment at steady-state using Eq. (S18) to obtain 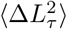, and 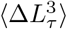 Using first and second central moments, we define the variance as 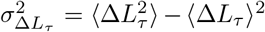. The closed form expressions of mean, variance are not reported here due to their complex expressions.

#### 4.2 Simultaneous binding model

As described in the main text, in this model, the filament undergoes elongation from states B and BF with rates *r*_1_ and *r*_2_, respectively. To write the master equation according to Eq. (1) described in the main text, we define the following matrices,

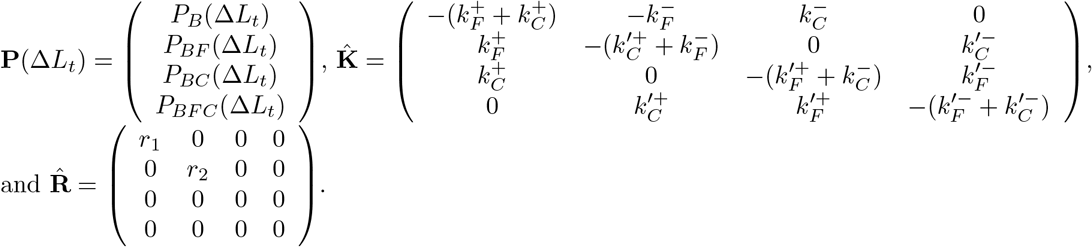

Using Eq. (S13), we derive the corresponding moments ⟨Δ*L*_*t*_⟩, and 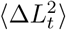 Using the first and second moments,the variance can be written as 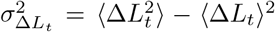. The closed form expressions of mean, variance are too cumbersome to be reported here.

Next, we calculate the mean, variance, and third central moment at steady-state using Eq. (S18) to obtain ⟨Δ*L*_*τ*_ ⟩, and 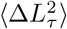. Using first and second central moments, we define the variance as 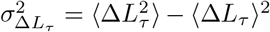 The closed form expressions of mean, variance are too cumbersome to be reported here.

